# A Unit Pipe Pneumatic model to simulate gas kinetics during measurements of embolism in excised angiosperm xylem

**DOI:** 10.1101/2021.02.09.430450

**Authors:** Dongmei Yang, Luciano Pereira, Guoquan Peng, Rafael V. Ribeiro, Lucian Kaack, Steven Jansen, Melvin T. Tyree

**Affiliations:** College of Chemistry and Life Sciences, Zhejiang Normal University, Jinhua, Zhejiang, China; Laboratory of Crop Physiology, Department of Plant Biology, Institute of Biology, University of Campinas (UNICAMP), Campinas SP, Brazil; Institute of Systematic Botany and Ecology, Ulm University, Ulm, Germany

**Author notes:** Corresponding Authors: Guoquan Peng, Melvin T. Tyree, or 1-802-777-3094.

**Keywords:** unit pipe, pneumatic model, gas diffusion, Pneumatron, vulnerability curves, embolism, xylem conduits, angiosperms

## Abstract

- The Pneumatic method has been introduced to quantify embolism resistance in plant xylem of various organs. Despite striking similarity in vulnerability curves between the Pneumatic and hydraulic methods, a modeling approach is highly needed to demonstrate that xylem embolism resistance can be accurately quantified based on gas diffusion kinetics.
- A Unit Pipe Pneumatic (UPPn) model was developed to estimate gas diffusion from intact conduits, which were axially interconnected by interconduit pit membranes. The physical laws used included Fick’s law for diffusion, Henry’s law for gas concentration partitioning between liquid and gas phases at equilibrium, and the ideal gas law.
- The UPPn model showed that 91% of the extracted gas came from the first two series of embolized, intact conduits, and only 9% from the aqueous phase after 15 s of simulation. Embolism resistance measured with a Pneumatic apparatus was systematically overestimated by 2 to 17%, corresponding to a typical measuring error of 0.11 MPa for *P*_50_ (the water potential equivalent to 50% of the maximum amount of gas extracted).
- Because results from the UPPn model are supported by experimental evidence, there is a good theoretical and experimental basis for applying the pneumatic method to research on embolism resistance of angiosperms.

## Introduction

According to the cohesion tension theory (CTT), water is transported in land plants under tensile conditions. The concept of tensile properties of water in vessels and tracheids of xylem tissue is contrary to the normal indoctrination of students of applied physics (e.g., mechanical engineering) because they are taught that only solids possess tensile properties, while liquids by “definition” do not. However, a pulling force can be applied to liquids enclosed in certain liquid containers to stretch and break them. CTT has withstood challenges over time (e.g., Benkert *et al*., 1995) through rebuttals based on reviews of past literature (Tyree, 1997), cell pressure probe experiments (Wei *et al*., 1999), and centrifuge experiments (Cochard *et al*., 2005). Tensile properties arise in water when confined to xylem conduits, which are xylem lumina with nanoscale pores in their cell walls. Therefore, the transport system from fine roots to the evaporative surface of leaves is composed of many intact pipes interconnected via pit membranes, which represent modified primary cell walls mainly composed of cellulose (Kaack *et al*., 2019).

Even though CTT has withstood the test of time, the transport of tensile water is prone to failure (Cochard *et al*., 2013). Immediately after a cavitation event (tensile failure), the conduit fills with a low-pressure void that consists primarily of water vapor. These voids eventually fill up with air at atmospheric pressure following Henry’s law, which describes the nature of the equilibrium between atmospheric gases and gases dissolved in water. One version of Henry’s Law can be written as:

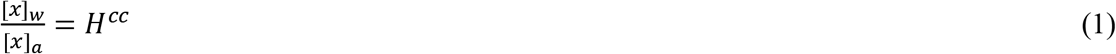

where [*x*] represents the mean concentration (mol L^−1^) of gas *x* in water, *w*, or air, *a*, depending on the subscript. *H^cc^* is a constant and approximately 10^−2^ for different gas species. The concentration of gas in the air phase comes from the ideal gas law:

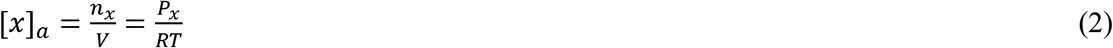

where *n_x_* is the number of moles of gas *x* in volume *V, P_x_* is the partial pressure of gas *x*, and *RT* is the universal gas constant multiplied by temperature in Kelvin.

In xylem conduits of plants, gases can appear after cavitation events if water remains in the tensile state and disappear if water returns to a non-tensile state slightly below atmospheric pressure. Modeling and experiments have determined the kinetics of bubble disappearance in conduits when the fluid pressure is near or above atmospheric pressure (Yang & Tyree, 1992; Tyree & Yang, 1992). Recent work has also focused on how long it takes a newly cavitated conduit to fully embolize, that is, when Ar, O2, and N2 reach partial pressures in the conduit equal to those in ambient air. Answers have come from theoretical models and experiments in which the focus has been on the speed of radial gas diffusion between conduits inside stems to the outside surface of the bark (Wang *et al*., 2015). This radial movement of gas from bubbles in conduits to the ambient atmosphere is basically controlled by Fick’s law of diffusion expressed in radial coordinates. This rate of diffusion of gases in water is very slow. Hence, even for stems less than 10 mm in diameter, the time for equilibrium can be hours to days depending on the diameter and the diffusion coefficient of gases in wet woody stems. In all of the studies cited above, axial diffusion was not included in the modeling, and the experimental designs for model verification inhibited most of the axial diffusion.

While these former models are useful and interesting, they do not apply to the new experimental situation inherent in the Pneumatic method (Pereira *et al*., 2016; Zhang *et al*., 2018) and the invention of the Pneumatron (Pereira *et al*., 2020; Jansen et al., 2020). The Pneumatron consists of an air pressure sensor connected by tubing to the cut end of a shoot with leaves, and it is used to measure the kinetics of diffusion from newly embolized, intact vessels to the embolized vessels at the cut surface of a terminal branch. This process involves axial diffusion of gases via hydrated pit membranes, which are typically only 0.2 to 1.3 μm thick (Li *et al*., 2016; Kaack *et al*., 2019). Over these short distances, diffusion can be quite quick. In axial transport, the median time, *t_m_*, for a gas to diffuse across a distance *s* is given by:

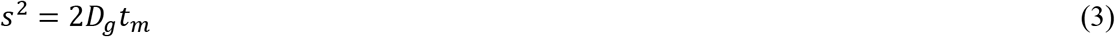

where *D_g_* is the coefficient of diffusion of the gas species *g* (in m^2^ s^−1^), and *s* is the distance (in m) that half the molecules traverse in time *t_m_* (in seconds). Gases in water have *D_g_* = 2 × 10^−9^ m^2^ s^−1^, so the time to diffuse through 1 μm of water is approximately 0.25 ms, but the time to diffuse through 0.01 m of water is 2.5 × 10^4^ s (≅ 7 h). Consequently, gases spread axially down cut branches more rapidly than radially through stems.

The Pneumatic method involves measuring the kinetics of gas movement down the axis of a stem while independently measuring the stem water potential with a stem hygrometer (thermocouple psychrometer connected to a stem) or by measuring the balance pressure of excised leaves with a pressure chamber. Since 2016, the Pneumatic method has been used to estimate the vulnerability curves (VCs) of woody species where the interpretation of what is measured is based on qualitative arguments, or by comparing pneumatic VCs to VCs measured by more conventional hydraulic (Pereira *et al*., 2016, 2020, 2021; Zhang *et al*., 2018, Sergent *et al*., 2020; Chen *et al*., 2020) and non-hydraulic techniques (Sergent *et al*., 2020; Chen *et al*., 2020; Guan *et al*., 2021).

What is clearly lacking in our full understanding of pneumatic measurements is a modeling approach of the gas diffusion kinetics along an axial pathway of conduits and radial pathways of stems. Therefore, the purpose of this study is to provide a theoretical background for pneumatic measurements, with the aim of proving that the theoretical kinetics of pressure change via diffusion of gases axially through intervessel pit membranes follows the proportion of embolized vascular tissue, either on a volume basis or hydraulic conductance basis. The model presented in this paper complements experimental evidence in recent papers (Pereira *et al*., 2020, 2021; Guan *et al*., 2021; Paligi *et al*., submitted).

## Model description

### Basic concepts of pneumatic measurements

In typical pneumatic experiments, the air pressure in all embolized vessels is assumed to be in equilibrium with the ambient air. In a measuring cycle, a partial vacuum (40 kPa absolute pressure) is drawn in <1 s at the cut surface of the stem. The stem is connected to a small, known volume of tubing, and air space is connected to a pressure transducer. Then, the vacuum pump is turned off, and the pressure is monitored every 0.5 s for ≤30 s, during which time the pressure increases by 10 to 20 kPa to an absolute pressure of 50 to 60 kPa. This is called air discharge. At the end of a measuring cycle, the air pressure is returned to atmospheric pressure for nearly 20 min to ensure that atmospheric pressure has been restored in all embolized vessels that are in axial contact with the cut open vessels.

### Basic idea of the Unit Pipe Pneumatic model (UPPn model)

All the theoretical calculations involve Fick’s law for the diffusion of gases, Henry’s law for gas concentrations at air/water interfaces, and the ideal gas law to relate air pressure to gas concentrations. Fick’s first law is used in radial or Cartesian coordinates as needed (Crank, 1975).

The Unit Pipe Pneumatic model (UPPn model) approximates the three-dimensional vessel network in angiosperm xylem (Zimmermann & Tomlinson, 1966). This model may provide a valid approach because the kinetics of axial gas exchange between embolized vessels (pipes) is much faster than that of radial diffusion between the surface of stems and embolized vessels. The model simulates the rate of gas extraction axially from embolized vessel lumina to the pressure transducer, and radially from the gas dissolved in the water of the surrounding xylem tissue. Axial transfer of gas occurs between closed (intact) vessels and cut-open vessels via diffusion through water spaces in the cellulose of intervessel pit membranes (Fig.1a).

**Figure 1.**
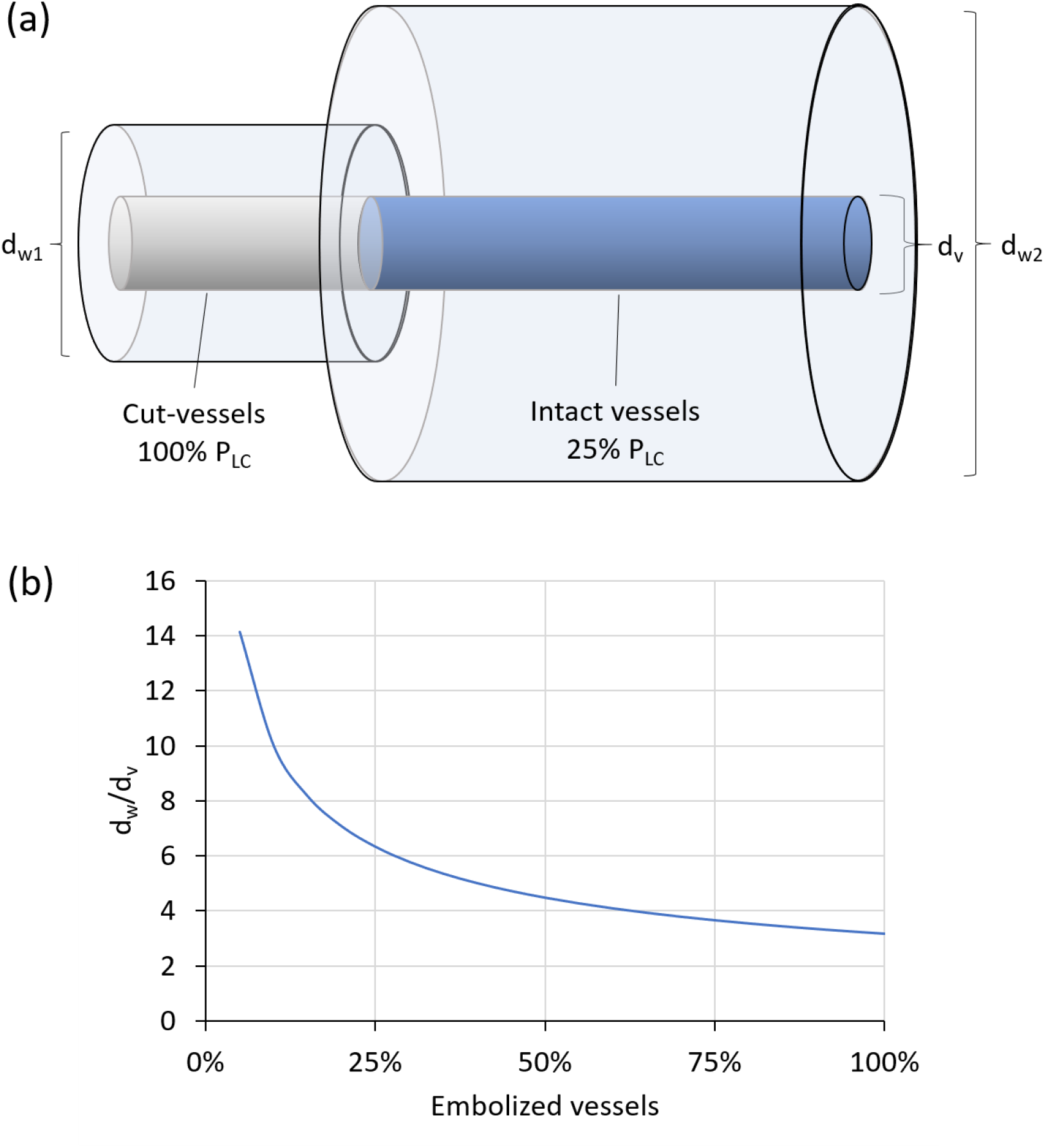
(a) A cut-open vessel and an intact vessel showing diameters of the vessel (far right, *d_w2_*), diameter of the cut vessel shell (far left, *d_w1_*), and diameter of the intact vessel shell in the middle (*d_v_*). (b) The ratio of wood to vessel diameters versus *PLC* used in the Unit Pipe Pneumatic model. The diameter of a wood volume is scaled so that all water-saturated wood is shared equally by the embolized unit pipes. Thus, if *PLC* is less than 100% in a region of wood, then the vessels that are embolized will share more wood diameter. Considering the fraction of wood area that is vessel lumina (*α_x_*), vessel diameter (*d_v_*), and wood diameter (*d_w_*) in the unit pipe model, the ratio *d_w_/d_v_* is given by (*α_x_ PLC*/100%)^0.5^, as shown in (b).

An important observation in the pneumatic method is that, at the beginning of an experiment, the quantity of gas discharge (measured in pressure change, *ΔP*) is minimal when there is zero embolism, but rises to a maximum difference when all vessels are embolized. If the vessel lumina hold only 10% of the water volume of woody stems, then the embolized vessels will contain approximately 5.8 times more moles of air than is dissolved in the 9-times-larger water volume surrounding the embolized vessels. Our model shows that the amount of gas extracted from the water of non-embolized wood is less than 10% of the gas extracted from the embolized vessels during the early part of the 30 s measuring cycle. This is because the rate of axial diffusion of gases from vessel to vessel is very fast and is rate-limited only by diffusion through the wet pit membranes. In comparison, the radial rate of diffusion is over an average radial distance of approximately 30 to 100 μm and is consequently much slower. Hence, axial diffusion is up to 100 times faster than radial diffusion.

### Transport and equilibrium coefficients used in the UPPn model

Henry’s law constants and Fick’s law coefficient of diffusion in air and pure water are shown in Table 1. In our model, we used a weighted average for Henry’s constant and the diffusion coefficient of air, *H^cc^* = 1.83 × 10^−2^ and *D_air,aq_* = 2.06 × 10^−9^ m^2^ s^−1^, respectively. Fresh (i.e., non-shrunken, non-dried) pit membranes have cellulose fibers with no lignin and an estimated pore volume fraction of 80%, which means that 80% of wet pit membranes are water (Zhang *et al*., 2020). The conventional method of dealing with a mixture of solids and water is to reduce the diffusion coefficient by the percentage of space that is water: 80%. The rate of oxygen diffusion in lignified wood of several species has been found to be one to two orders of magnitude less than that in pure water (Sorz & Hietz, 2006), so the value for radial diffusion should be reduced accordingly. The coefficient of diffusion of gases in air is four orders of magnitude larger than that in water, and in embolized vessels, the mass flow of gases can accelerate pressure equilibrium even more.

**Table 1.** Henry’s law constants (*H^cc^*), diffusion coefficients in water (*D_g,aq_*) and air (*D_g,air_*), and gas concentration in water (*C_aq_*) and air (*C_air_*). RT = 24.8 L bar mol^−1^, 1.013 bar = 1 atm. Avg = weighted average = sum of %air × D_g,aq_ (or H^cc^). If Ar is ignored, the weighted averages are slightly different.

| Gas | $D_{g,aq}$<br>(m<sup>2</sup> s<sup>-1</sup>) | $D_{g,air}$<br>(m<sup>2</sup> s<sup>-1</sup>) | $H^{cc}$<br>($C_{aq}/C_{air}$) | $C_{aq}$<br>(mM) | $C_{air}$<br>(mM) | Air<br>(%) |
| --- | --- | --- | --- | --- | --- | --- |
| Ar | 2.00E-09 | 7.03E-05 | 3.43E-02 | 1.40E-02 | 0.41 | 1 |
| O<sub>2</sub> | 1.88E-09 | 1.58E-05 | 3.18E-02 | 2.60E-01 | 8.17 | 20 |
| N<sub>2</sub> | 2.10E-09 | 1.76E-05 | 1.49E-02 | 4.81E-01 | 32.27 | 79 |
| Avg | 2.06E-09 |  | 1.83E-02 |  |  |  |

### The UPPn model

The UPPn model was used in this study because it is simple enough to be programmed and solved in an Excel spreadsheet. The Excel spreadsheet can be created with all equations embedded into cells and solved in a printed mathematical format. The Excel spreadsheet also has most of the intermediate and all of the final calculation results presented automatically in graphical format. This spreadsheet is available in the Supplementary Information (Table S1). Hence, only the essential features of the model are introduced below.

In UPPn, we modeled for the two slowest processes: (1) axial gas diffusion, rate-limited by diffusion across pit membranes, and (2) radial gas diffusion in concentric rings surrounding a single, embolized vessel. The radial path length and volume were adjusted in each calculation so that all wood volume in a stem was divided equally between all embolized vessels. Hence, if a model stem had *V_s_* volume of non-embolized space (vascular and non-vascular volume) and a count of *N_e_* embolized vessels each of volume *V_v_*, then each vessel was assumed to be surrounded by *V_s_/N_e_ - V_v_* of non-embolized wood volume per vessel. Therefore, the water-saturated wood volume surrounding a unit embolized vessel changed with percent embolized vessels. As shown in Fig. 1a, the unit pipe consisted of a cut-open vessel (left; colored gray) and one or more intact vessels (right; colored blue). Every cut-open vessel on the left was embolized and hence had the minimum amount of water-saturated wood in the outside radius (white). Assuming 25% embolism in Fig. 1a, the woody tissue volume on the right side was twice the diameter and four times the volume as on the left side. As the percentage loss of conductivity (*PLC*) varied, the ratio of water-saturated wood to embolized vessel diameter changed accordingly (Fig. 1b).

The UPPn model provides an adequate resolution of the time course of pressure changes in vessels after a partial vacuum is drawn on all cut-open vessels. Because stem samples prepared for pneumatic measurements are cut in the air and the cut-open vessels become quickly embolized, they function as an extension of the discharge tubing. We assumed that only mature functional vessels were capable of forming embolisms. Therefore, living immature vessels, cambium, and living bark cells have no embolism. Hence, unit pipes near the boundary between mature and developing vessels would also receive some air by diffusion through the water-saturated wood volume, cambium layer, and living phloem in bark. The UPPn model slightly underestimated the rate of pressurization of vessels near the surface of the stem, but because radial diffusion is much slower than axial diffusion, this amount of error was acceptable over the time domain of the model (typically 15 s). The fastest diffusion occurred axially through the intervessel pit membranes over an average length of 3 to 30 cm. Therefore, most of the air extracted by the Pneumatron came from axial diffusion because of the high axial diffusional rates, and because the amount of air in the aqueous solution was only 2% of the concentration in the embolized vessels.

Conventionally, Pneumatron data is used to compute the ratio of gas discharge after a fixed time of 15 to 30 s. Empirically, the amount of gas discharged into a fixed volume *V_o_* causes a pressure increase by Δ*P*. The pressure discharge is the least, Δ*P_min_*, at the start of an experiment and greatest, Δ*P_max_*, at the end. A dimensionless value is calculated that correlates with the percentage embolism:

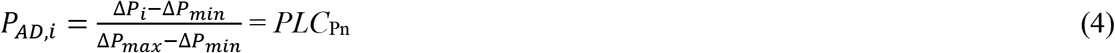

where *P_AD,i_* is the i^th^ percentage air discharge but could also be called *PLC*_Pn_, the assumed measured *PLC* by the Pneumatron.

The same relative discharge is calculated for ideal gases whether one uses pressure, concentration, or moles of gas in Eq. (4). The assumption is that *P_AD,i_* is closely related to *PLC_Pn_*, which could equally be percent embolized vessel volume or percent loss of hydraulic conductivity in a unit pipe model. The purpose of these models is to investigate how the theoretically computed *P_AD,i_* relates to hydraulic *PLC* values. In the UPPn model, all vessel diameters are equal. When there is a range of vessel diameters (*d_v_*), and if some of the diameters are more vulnerable than others, then the percent vessel volume is proportional to 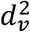, but the conductance is proportional to 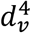.

### Radial and axial diffusion through the wood

In the numerical simulation, there are only two types of rate constants that must be calculated, and they depend on the values listed in Table 2. One type of rate constant is for radial diffusion, the other is for axial diffusion, and both use Fick’s first law for diffusion in radial or Cartesian coordinates (Crank, 1975).

**Table 2:**
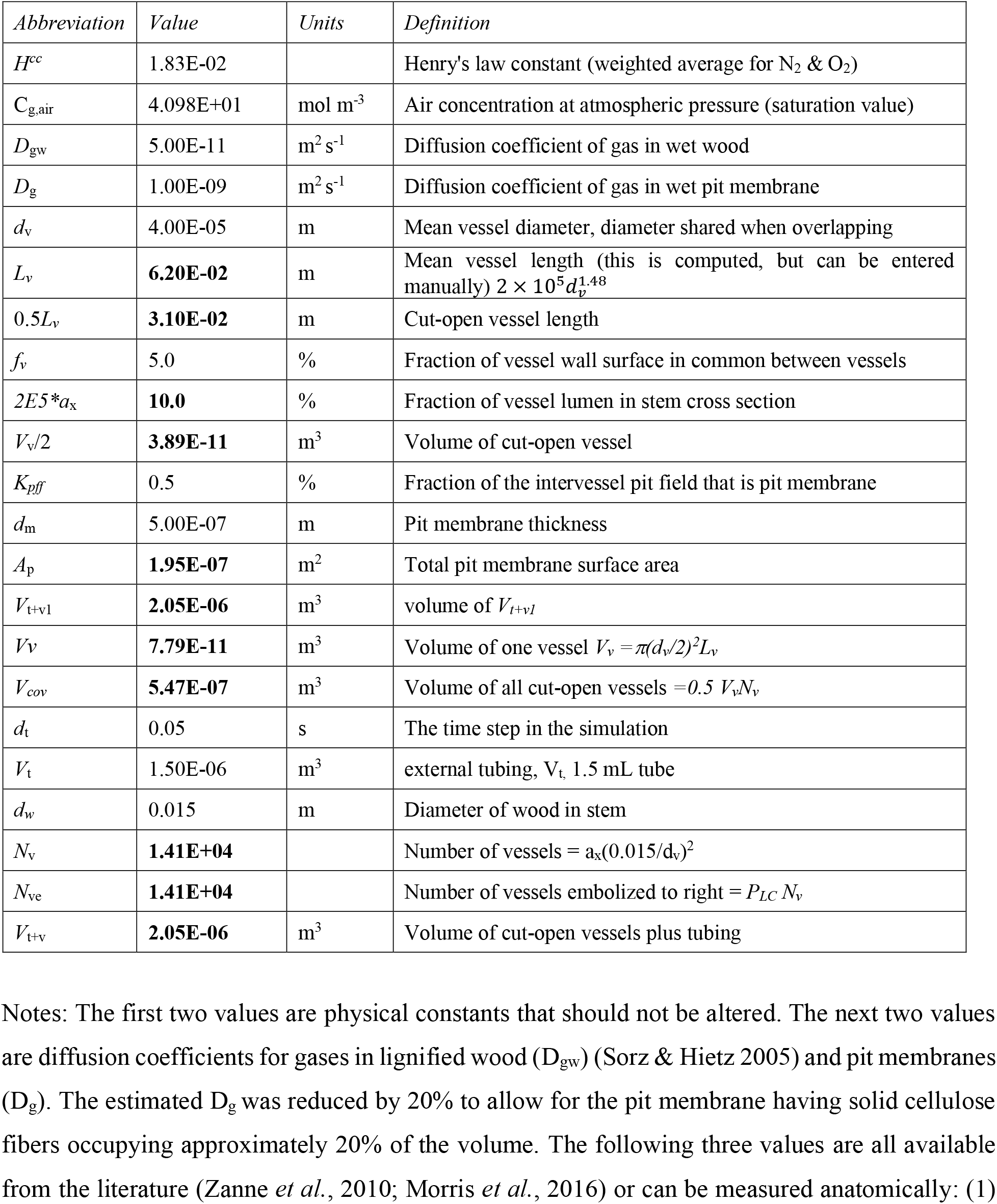
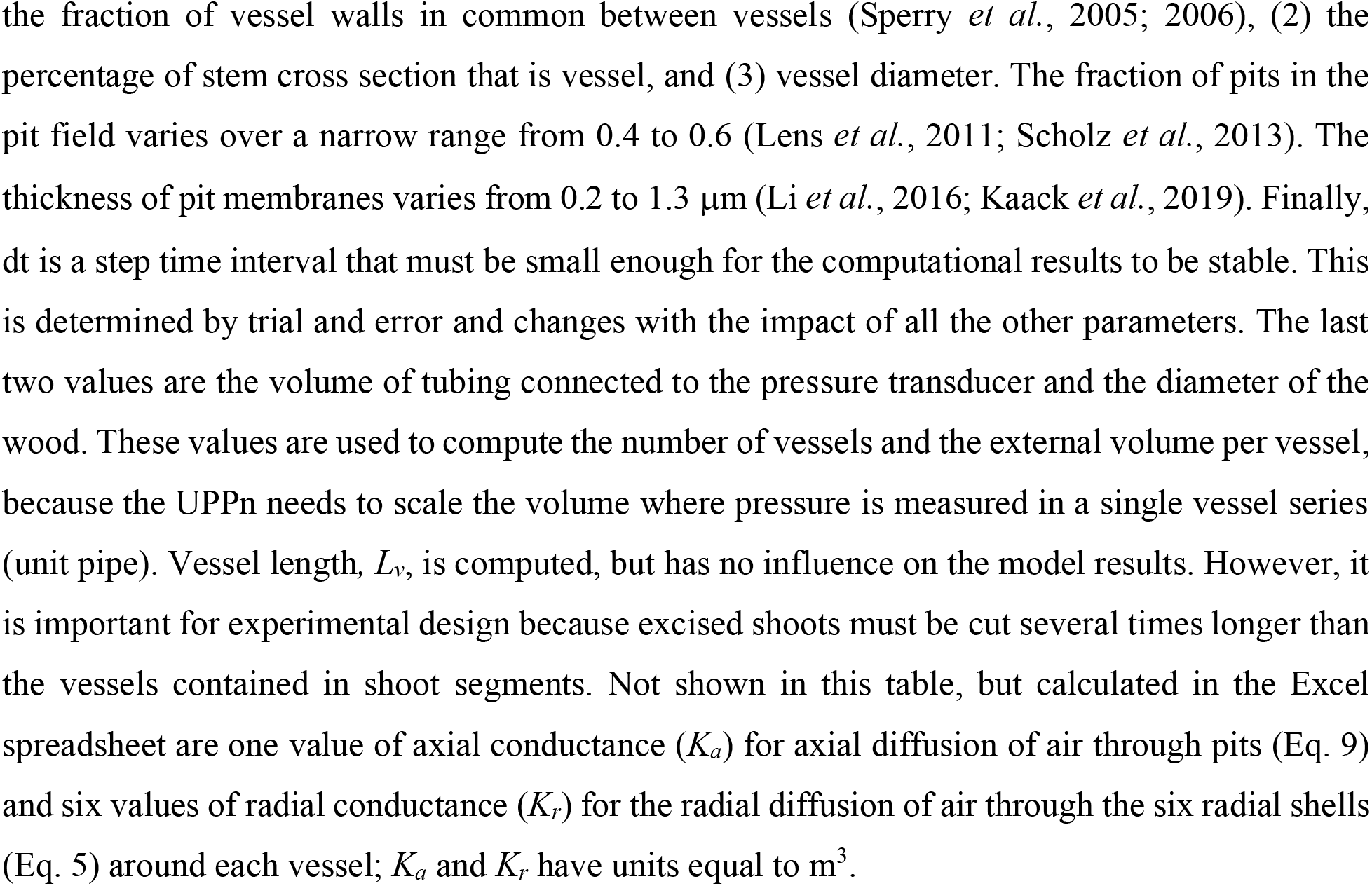
Values that must be entered in the UPPn model Excel spreadsheet. The non-bold values are entered by the user; the **bold** values are calculated by Excel.

The radial diffusion pathway was divided into six concentric rings of radial step *Δr*. The rate constant for each concentric ring was unique, depending on the radial diffusion coefficient of gases in lignified wood (*D_gw_*), vessel length (*ΔL_v_*), radius of the i^th^ ring (*r_i_*), volume of each cylindrical concentric ring (*V*), and time step (*Δt*). The Excel spreadsheet computes the change in concentration of gases (*ΔC*) in time increment (*Δt*). In the radial and axial paths, the rate constants for diffusion had *Δt* included. This is done to increase the speed of update of the *ΔC* values in each row of the Excel spreadsheet. The meaning of all symbols is given in Table 2. The rate constant for radial diffusion is:

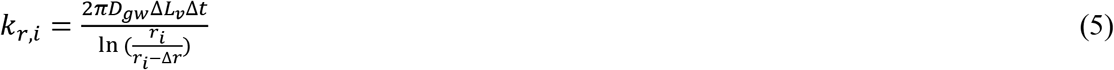

The equation for the last (n^th^ = 6^th^) ring is:

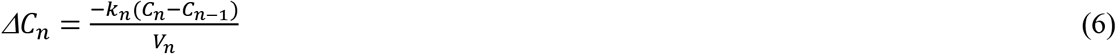

For the intermediate i^th^ ring, the equation is:

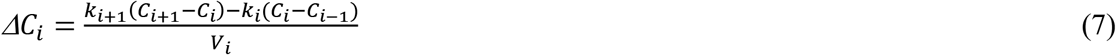

For the innermost ring next to the vessel, the equation is:

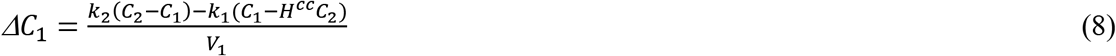

Regarding axial diffusion, the rate constant of each vessel in series is:

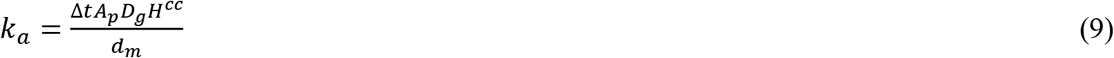

where *A_p_* and *d_m_* are the area and thickness, respectively, of the pit membranes in the axial path.

Then, the change of concentration in each time step is:

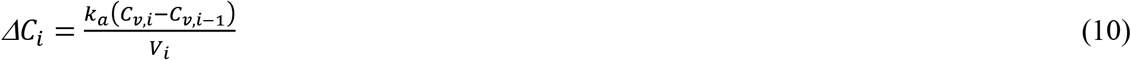

All of the above equations are included in the appropriate Excel cells (Table S1). At time zero, the intact vessel is assumed to be filled with air at atmospheric pressure, but the cut-open vessel has been reduced to an initial pressure (concentration) given by the ideal gas law. The Pneumatron draws down the pressure over a period of 1 s or less. Then, after each time step, Eqs 5, 7, 9, and 10 are used to calculate the change in concentration in each Excel cell where each cell represents a location in the vessel or surrounding water-filled wood. Then, before the next time step, each concentration is adjusted by the corresponding *ΔC* value computed for the time interval *Δt*. The answers are put into the Excel row corresponding to the elapsed time. The reader is referred to supplemental material for the Excel spreadsheet and a description of the basic layout of the sheet (Table S1). Some of the key anatomical values (Table 2) that are needed in the Excel spreadsheet are the diffusion coefficients in wood (Sorz & Hietz, 2006), vessel dimensions in wood (Zanne *et al*., 2010; Morris *et al*., 2016), and pit membrane thicknesses in vessels (Li *et al*., 2016; Kaack *et al*, 2019).

## Results

### Pressure dynamics over a 150-s period

First, consider the case for 50% of vessels embolized. A UPPn model simulation of pressure is shown in Fig. 2. The only parameter measured by the Pneumatron is the pressure in the cut open vessels (blue line, labeled ‘#0’) computed from the concentration (*n/V*) by the ideal gas law *P = (n/V)RT*. The pressure in the first intact vessel started out at atmospheric pressure. The temporal dynamics of the axial pressures simulated in ten vessels connected axially are shown in Fig. 2. It can be seen that, in the first 15 s, almost no change in pressure was registered beyond the fifth vessel in series. The conclusion is that the Pneumatron can detect the pneumatic influence of only the first few vessels in the axial chain of embolized vessels.

**Figure 2.**
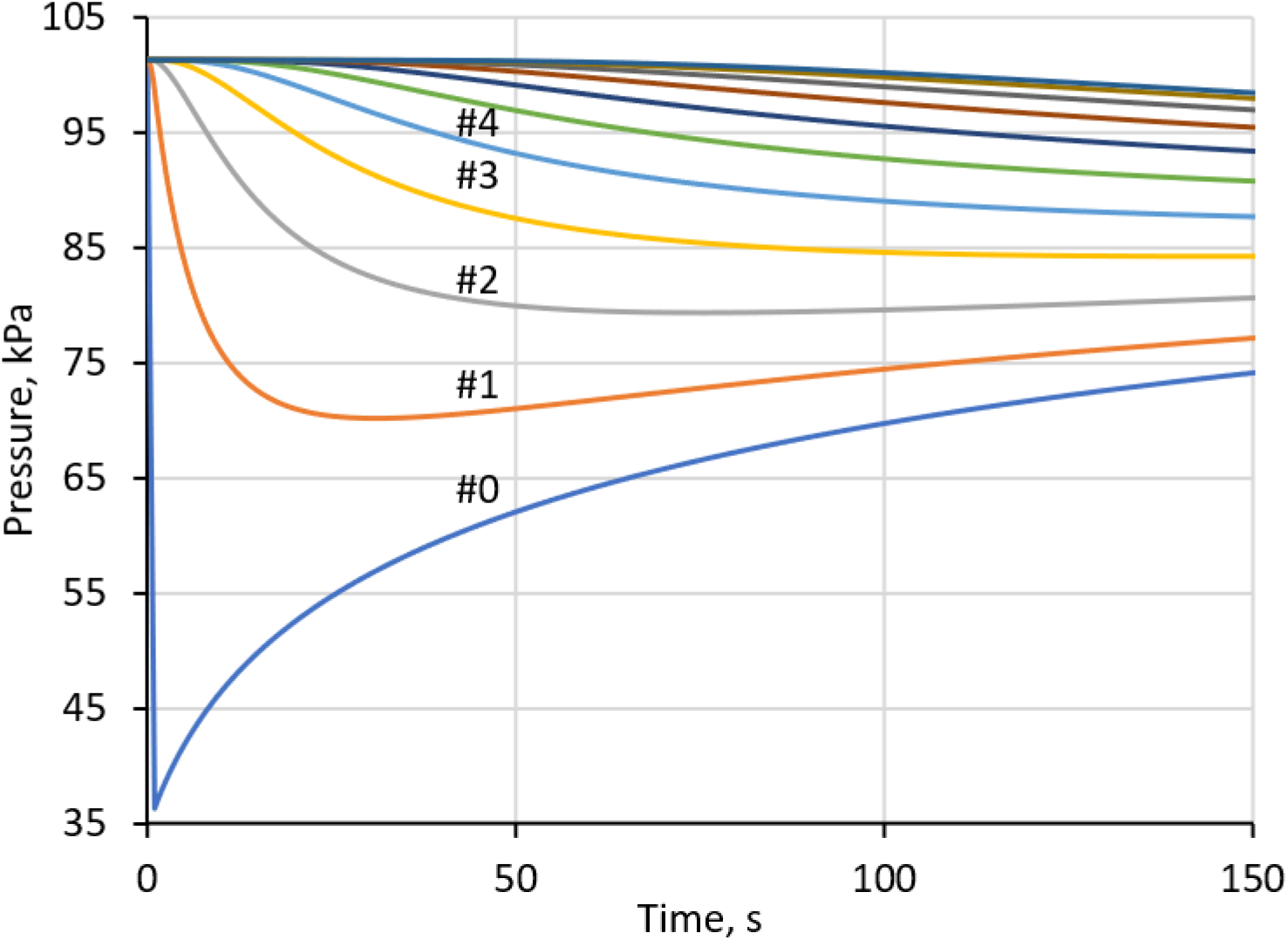
Simulated absolute pressures according to the Unit Pipe Pneumatic model measured by the pressure transducer (blue, #0) and as computed in the intact, embolized vessel. All vessels start out at full atmospheric pressure. In the first 15 s after drawing a partial vacuum, almost no change is detected in the seventh vessel down the chain. The pressure transducer is in pressure equilibrium with the cut vessels. Vessel number is indicated below each line where space permits.

According to the UPPn model, most of the extracted gas came from the embolized vessels rather than the aqueous phase surrounding the vessels. After 15 s, the total gas drawn into the volume space connected to the pressure transducer (i.e., the discharge tube) came from the embolized vessel space and the aqueous phase surrounding the embolized vessel. Based on Fick’s law of diffusion, the total amount of gas extracted was 1.53 × 10^−9^ moles, of which 91% came from the embolized space and only 9% from the aqueous phase. Even after 150 s, the percentages of gas extracted from the gas and liquid phases were 84% and 16%, respectively.

Each solution depends on the parameters shown in Table 2 and the assumed percentage of embolized vessels. In the solution above, the model assumed 50% embolism. Hence, the radius of the external tissue was minimum in the cut-open vessels and greater in the 50% embolized zone. The radial gas concentrations in the six concentric rings of water-filled tissues to the right of the cut-open vessels were computed (Figs. 1a and 3a). The concentric ring concentrations of gas reached a minimum value and then began to rise again, which was a consequence of the nature of the simulation. The concentrations of gas in the concentric rings of intact vessels changed less than in those of the cut-open vessels. After >50 s, the gas concentrations in the rings started to increase because of air-entry from vessels further down the chain. The third and sixth vessels along the axis showed much smaller changes in dissolved gas concentration over the same 150 s (Fig. 3b, c). The conclusion from this simulation is that, even if there are many vessels in an axial chain that are embolized, the amount of gas extracted in the first 15 s of the extraction process comes mostly from the first two intact vessels.

**Figure 3.**
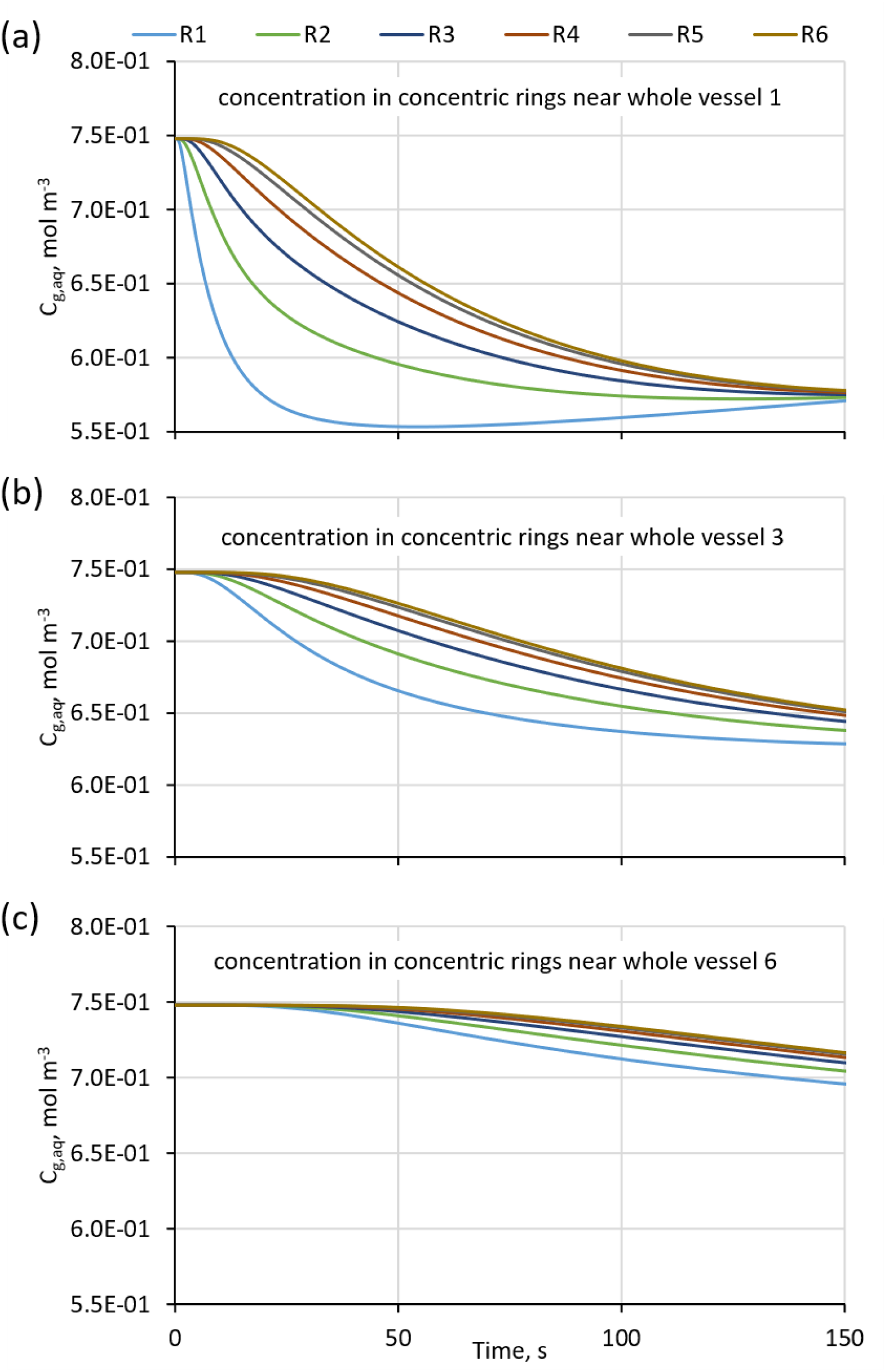
Temporal dynamics of the axial concentrations of air in the aqueous phase of water-filled walls and wood fiber cells when 50% of the vessels are embolized. Each colored line equals the concentration of gas in water in a concentric ring of wood around the vessel. In the legend, R1, means the ring nearest the first vessel, R2, means the second ring from the vessel, etc.

### Pressure kinetics with zero embolism (P_min_)

In this case, the pneumatic model solves for a cut-open vessel that is half the mean vessel length. There is no axial draw of air because all axial intact vessels are water-filled. Therefore, we must consider only the exchange of air from water-filled wood radially adjacent to the cut-open unit pipe. In the Excel spreadsheet (Table S1), this is easily programmed by changing a Boolean variable (cell L44) from false to true, which indicates that the spreadsheet does not consider axial flow from vessels distal to the cut-open vessels because there are no embolized, intact vessels to allow axial flow (except for the cut-open vessels). The spreadsheet for this simple case computed a very small change in moles of gas in the external chamber connected to the pressure transducer: 7.45 × 10^−11^ mol after 15 s and 1.54 × 10^−10^ mol after 150 s. Theoretically, this should be the same as setting *PLC* (cell O39) to zero, but this will cause the program to crash because this value is used to compute the radial distance of water-filled tissue between equally spaced embolized vessels. In practice, it is better to maintain the *PLC* above 5% even though the Excel spreadsheet computes outputs for values as low as <0.1%, but the output grows increasingly inaccurate for small values of *PLC* in cell O39.

### Pneumatron absolute pressure curves and PLC_Pn_ curves versus model input PLC

The impact of the extractable gas (model output) as the input PLC increases from zero (cell L44 = true) to 100% (cell L44 = false, cell O39 set to the desired *PLC*) is shown in Fig. 4a. Minimal change in pressure Δ*P_min_* occurs at 0% *PLC* input, and the maximum *ΔP*_max_ occurs when 100% *PLC* is inputted. Readers should remember that, in the UPPn model, an increase in *PLC* is equivalent to a decrease in the radius of non-embolized water/tissue around the embolized vessel. The conventional method of estimating the *PLC* in previous experimental studies was to pick some time interval (e.g., 15 or 30 s) and then compute *PLC_Pn_* following Eq. (4). Taking the Δ*P_i_* measurements at 15 s for every curve in Fig. 4a produces the results shown in Fig. 4b. The relationship is slightly curvilinear, with a maximum deviation of approximately 5% at the 50% *PLC* input value.

**Figure 4.**
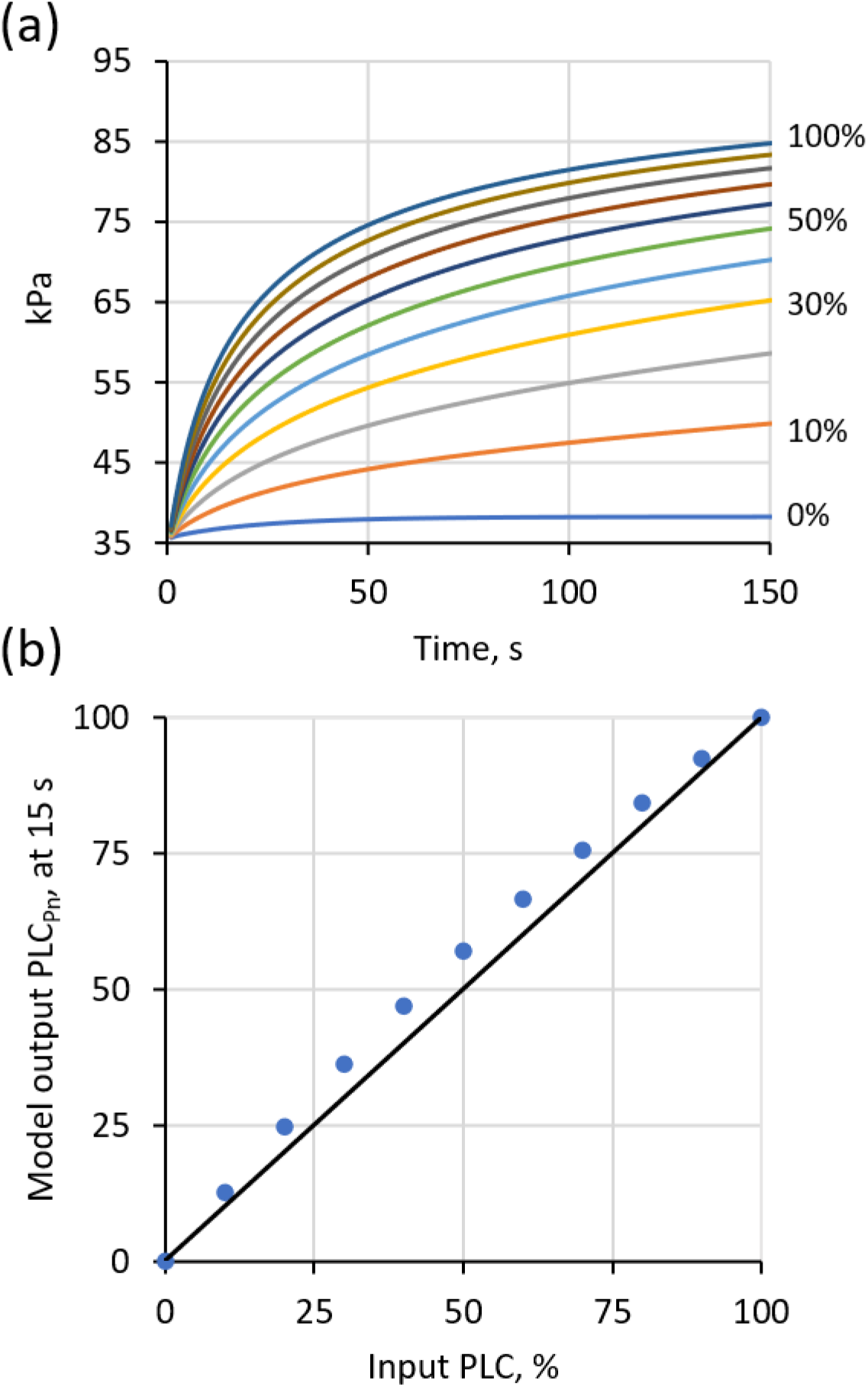
(a) Temporal dynamics of simulated pressure output (kPa) at the pressure transducer with varying *PLC*. In the simulation, the partial vacuum is drawn in the 1-s interval before time 0 on the x-axis. The different curves give the gas pressure change when varying the percentage of embolism in the stem (shown on the right). (b) A plot of simulated *PLC* from the model versus the input *PLC*. The points are taken from the curves in (a) at t = 15 s using Eq. (4). The black line represents the 1:1 line.

### Sensitivity analysis

The rate-limiting steps for gas movement occurred when gas was forced to move through water by diffusion. When there was no embolism at all in intact vessels, the gas extraction was limited to what came out of solution from the surrounding water radially connected to the cut-open vessel [Δ*P_min_* in Eq. (4)]. When vessels were embolized, the maximum air extraction occurred from the embolized vessel plus the extraction from the water-filled tissue surrounding the vessel [Δ*P_max_* in Eq. (4)]. There would be a perfect match between the volume of embolized intact vessels and *PLC_Pn_* in Eq. (4) if no gas was drawn out radially from the surrounding tissue.

The rate of gas extraction from intact vessels was faster than that from surrounding water because the diffusional path length equaled the pit membrane thickness per vessel (0.2 to 1.2 μm), whereas radial diffusion was over a distance of 60 to 150 times the typical pit membrane path length (30 μm to 75 μm depending on vessel diameter) when the volume fraction of the vessel occupied 10% of the cross-sectional area of the wood.

The mass flow rate of gas down the length of the vessel lumen was much more efficient than that of diffusion. The maximum gas flow down the axis of the vessel occurred in the first vessel adjacent to the cut-open vessel immediately after the vacuum was drawn. The theoretical pressure drop needed to maintain flow down the entire vessel length was calculated to be <10^−4^ Pa out of nearly 10^5^ Pa initial pressure; hence, pressure gradients through the length of intact vessels can be ignored compared with the pressure difference across intervessel pit membranes.

In most simulations, the amount of gas extracted in the first 15 s (the measuring cycle of the Pneumatron) was >90% from the vessels. Vessel length had no impact on the fraction of extracted gas because increasing vessel length increased the moles of air in the embolized vessel and increased in exact proportion to the moles of air in the surrounding water-saturated tissue. In contrast, increasing the vessel diameter increased the percentage of gas extracted from the embolized vessels, even though the volume of water-saturated annuli dramatically increased with the vessel diameter (Fig. 5). If gas extraction from embolized stems was 100% from vessel lumina with nothing from the water-saturated annuli around the vessels, then the Pneumatic technique would provide an accurate measurement of the volume of embolized vessels. However, a precision of 85 to 95% was not bad given how easy and inexpensive the Pneumatic method and Pneumatron respectively are. These percentages were computed for an analysis period of 15 s. The decision about the appropriate time interval for analysis of real experiments versus simulated data is addressed in the discussion with one experimental example. All the percentages in Fig. 5 were computed for the 50% embolism case. The agreement improves slightly as the input *PLC* decreases. For example, for a 40-μm diameter vessel, the percentage gas extraction increased from 90.3% at 100% PLC input to 91.0% at 10% embolism input.

**Figure 5.**
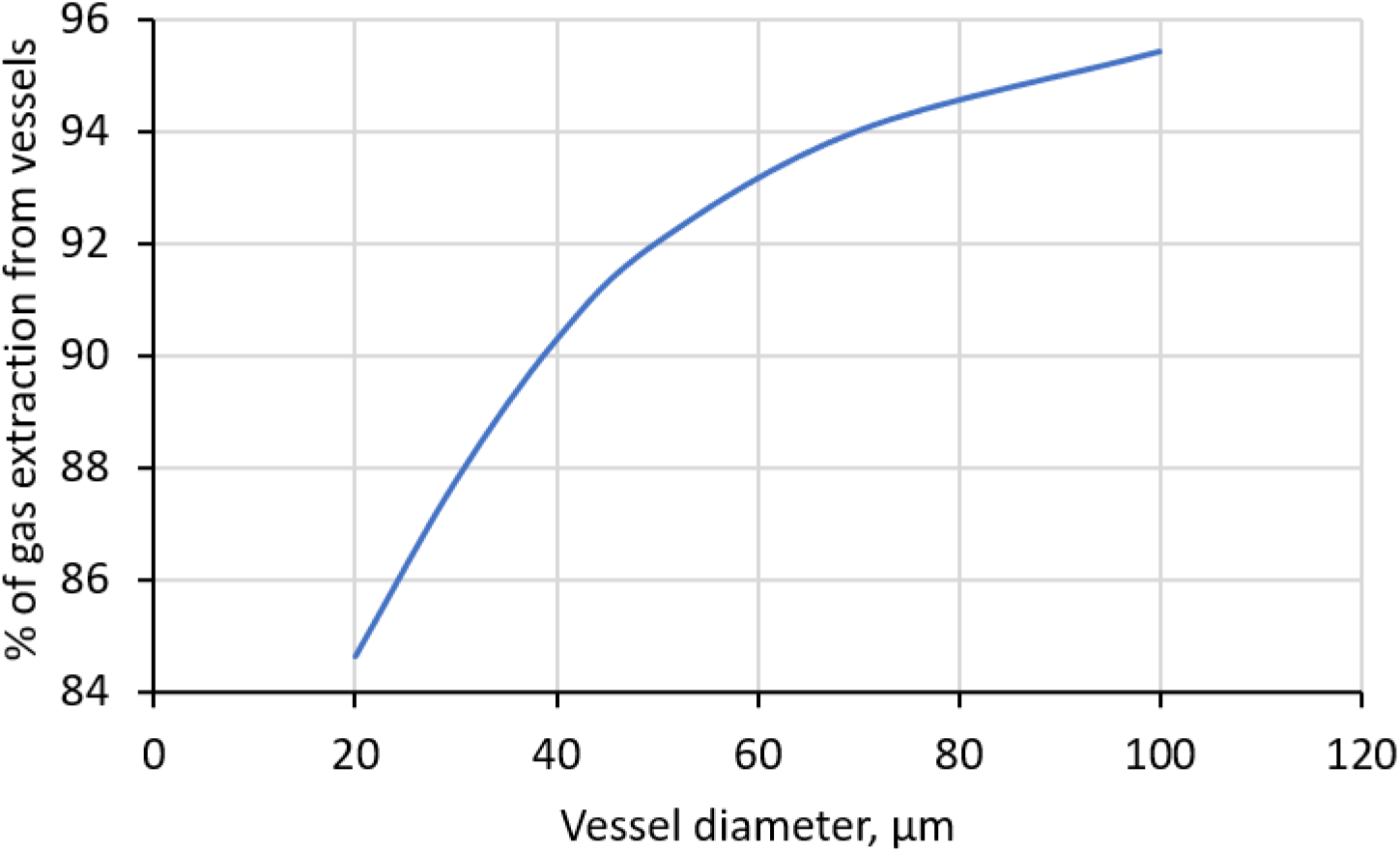
The percentage of gas extracted from the vessel in the first 15 s of the Unit Pipe Pneumatic model simulation as affected by vessel diameter. The percentages are percent of total gas extraction from the embolized vessels plus the air extracted from the water surrounding the embolized vessels.

So far, we have presented results from the UPPn model using only one set of input parameters, as shown in Table 2. These are the default values available to readers who wish to download our Excel file. The only factor that changed was the input *PLC* value. We must now examine how much the model output changed when other input values were selected that might have influenced the UPPn model. Based on quantitative anatomy, the values in Table 2 cannot be independently varied and expected to be meaningful (Sperry *et al*., 2005, 2006; Hacke *et al*., 2006). For example, vessel length (*L_v_*) scales with vessel diameter squared, and 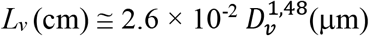 (Liu *et al*., 2018); converting both sides to meters yielded 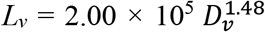. In addition, end wall resistivity in pits scales with lumen resistivity with a slope near one. The initial UPPn model took these factors into account when default values were loaded in Table 2 (copied from the Excel file available for download). All values of *L_v_* in Sperry *et al*. (2005, 2006) and Hacke *et al*. (2006) seem rather short compared with those in recent literature (Liu *et al*., 2018), but we can safely skip this debate because the UPPn model was insensitive to *L_v_*.

### Influence of axial conductance (k_a_) on Pneumatron results

The axial conductance (*k_a_*) used in the calculations is a diffusional conductance for air, whereas the statement about hydraulic resistivities being nearly equal refers to the resistance to water flow per unit length of wood. Pit membranes are typically 0.3 to 0.5 μm thick (range about 0.2 to 1.2 μm) compared with vessel lumen lengths that are 10^4^ to 10^6^ times longer (5 to 50 cm or more). The UPPn model considers the rate-limiting step of the diffusion of gases through the water-filled spaces of pits that are approximately 80% water by volume in fresh pit membranes (Zhang *et al*., 2020).

The model predicted, after 15 s, that >90% of the gases drawn into the tubing connected to the pressure transducer came from axial gas extracted from embolized vessels, and less than 10% came from gases dissolved in the water of fully hydrated tissue surrounding the vessel. Therefore, we started with the assumption that factors influencing *k_a_* dominated the gas-extraction process. There are four constants in Eq. (9). The value of *Δt* is the time step used for the iterative solution of the equations, and the only thing we do with that is pick a small enough value so that the solution is stable. The two most variable constants are the pit membrane area between adjacent vessels (*A_p_*) and the thickness of the pit membrane (*d_m_*). The value of *A_p_* seems to range over two orders of magnitude from 3 × 10^−7^ to 3 × 10^−9^ m^2^ (Wheeler *et al*., 2005; Hacke *et al*., 2006; Lens *et al*., 2011; Jansen *et al*., 2011; Scholz *et al*., 2013) and has been suggested to be negatively correlated with vulnerability to embolism (Wheeler *et al*., 2005; Hacke *et al*., 2006; but see Kaack *et al*., submitted). The value of *d_m_* was not considered in Hacke *et al*. (2006), but we now know it ranges from 0.2 to 1.2 μm and is positively correlated with the tension at 50% embolism (*T_50_*) (Li *et al*., 2016; Kaack *et al*., 2019). Since *k_a_* is a function of the ratio of *A_p_/d_m_*, it seemed reasonable to explore how this ratio changes over a factor of 10 from 2 to 0.2 m, and this was accomplished by changing *k_a_* from 5 × 10^−11^ to 5 × 10^−12^; *k_a_* is the diffusional conductance of gas in wet pit membranes times *Δt* (time step = 0.05 s). The results, pressure vs. time, are plotted in Fig. 6a, and all simulation curves were convex upward, as shown in Fig. 4b. Figure 6b is a plot of the simulated value of *PLC_Pn_*, that is, the model output value when the input value is 50% *PLC*.

**Figure 6.**
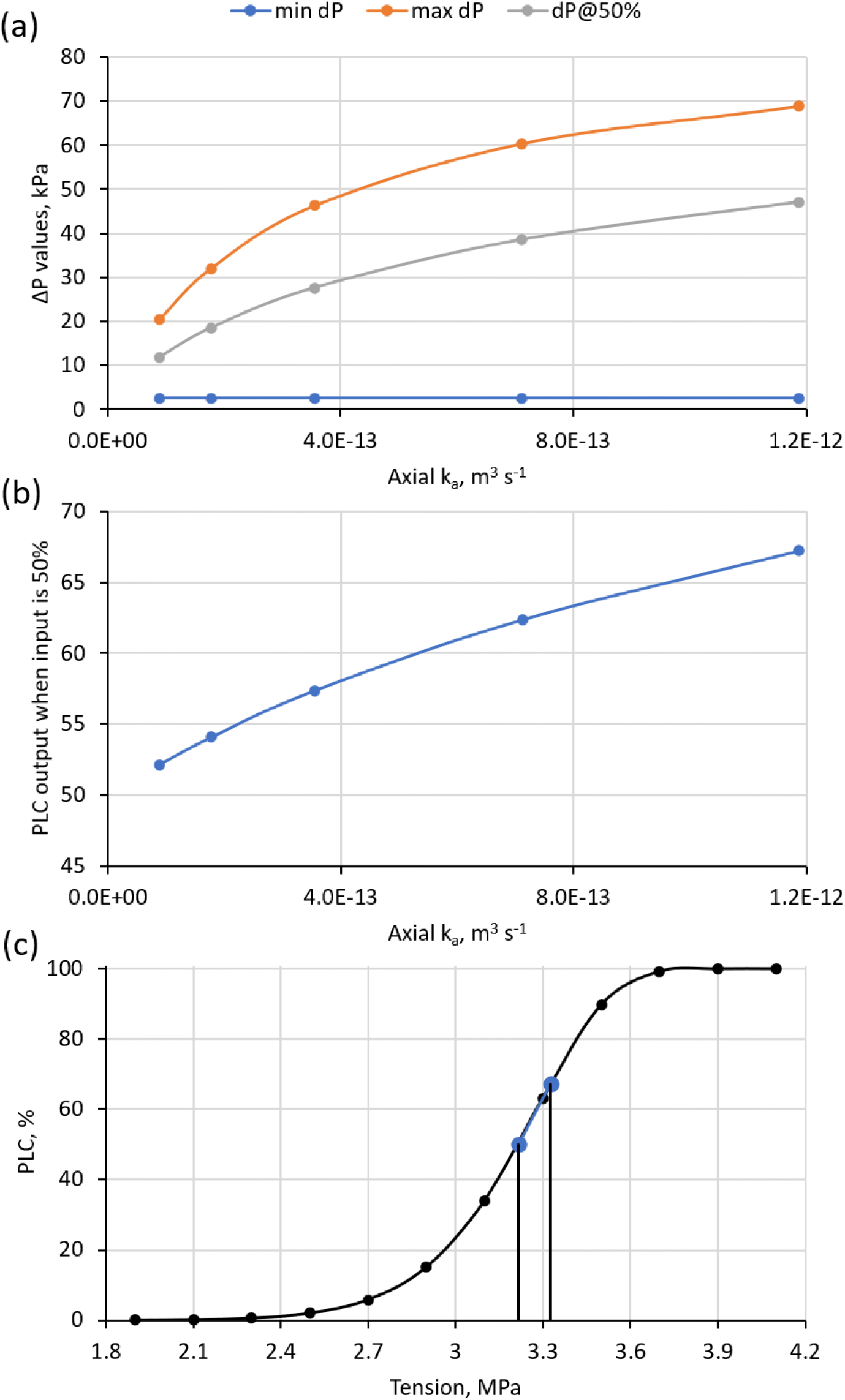
Theoretical impact of axial conductance of pit membranes to air diffusion (*k_a_*) on the computed change in gas pressure (*ΔP*). (a) The computed pressure values after 15 s for 0 (*ΔP*_min_), 50% (*ΔP*_50_), and 100% (*ΔP*_max_) embolism as affected by *k_a_*. (b) The pneumatic value of *PLC_50_* as affected by *k*_a_. (c) The maximum likely error in determining the tension at 50% *PLC* (*T_50_*) is shown on a typical vulnerability curve, assuming an overestimation of 17% in *PLC_50_*.

The simulation demonstrated that changing *k_a_* over likely values in plants increased the model-computed values of Δ*P_max_* and *ΔP50* when the real input value was 50% (Fig. 6a). Δ*P_min_* was unaffected by *k_a_* because there were no embolized vessels to deliver gases (Fig. 6a). The computed tension at *PLC_50_* from the Pneumatron was in error of 2% to 17% from the true value of 50% as *k_a_* increased (Fig. 6b), and hence, the Pneumatron always overestimated the tension of *PLC_50_*. However, such overestimation of *PLC_50_* would have a low impact on *T_50_* (≤0.11 MPa), as shown in a typical VC (Fig. 6c). This error magnitude appears to be acceptable and less than or equal to the typical disagreement in *T_50_* values measured by different hydraulic methods on the same species (see review by Cochard *et al*., 2013).

### The impact of radial diffusion of gases on Pneumatron results

When a vessel is embolized, gas can diffuse from the hydrated tissue immediately adjacent to the embolized vessel. When the Pneumatron pump is turned on, it draws a partial vacuum at the end of the cut stem, which causes axial diffusion of gases from the nearby embolized vessels, which are initially at atmospheric pressure. Then, as the intact vessel pressure drops, there is a tendency of gases to diffuse from the surrounding tissue. The model already demonstrated that the axial rate of diffusion was faster than the radial rate. This was the consequence of two factors: (1) the distance of radial diffusion was about 100 times greater than the axial diffusion distance in the water of pit membranes; and (2) the coefficient of diffusion was up to two orders of magnitude less in water-saturated wood (Sorz & Hietz, 2006) compared with that in pure water. The values of the O2 diffusion coefficient measured in water-saturated wood were 1 × 10^−11^ to 2 × 10^−10^ m^2^ s^−1^, which were lower than that in pure water (2 × 10^−9^ m^2^ s^−1^). The default value used for our calculations was 5 × 10^−11^. We used this low value because the wood samples studied by Sorz & Hietz (2005) were from stems that still had some embolism (5–20% gas volume). Exactly where this gas was located was not specified by the authors, but the range of gas contents was close to the percentage of stem volume that contained vessel lumina in most woody species (5–20%) (Zanne *et al*., 2010; Morris *et al*., 2016). Increasing the gas content by another 10% typically caused a ten-fold increase in the diffusion coefficient. This is because the pathway of gas movement involves water and air in parallel and series pathways, and the coefficient of diffusion of gas in gaseous medium is 10^4^ times larger than that in water (Table 1). Hence, the 5 × 10^−11^ value we used may still have been too large and was already five times the minimum value (Sorz & Hietz, 2006). Starting with the default values (Table 2), the percentage of gas extracted from radial pathways was 8.2% of the total amount of gas extracted. Decreasing the diffusion coefficient by a factor of two decreased this percentage to 5.4%, and increasing the diffusion coefficient by a factor of two increased the radial extraction to 12.2%. The readers are invited to enter their own values in the Pneumatron Excel spreadsheet. Hence, our sensitivity analysis suggests that *PLC_Pn_* may be a robust estimate of hydraulic *PLC*.

## Discussion

### The initial rate of gas extraction is the best predictor of embolism

Common sense, experimental data, and theoretical models all point to the most important conclusions: the initial rate of gas extraction is the best predictor of the cut shoot *PLC*. In the first second or less, gas is extracted only from the first intact, embolized vessels from the base of an excised organ. The gas pressure must drop in the first vessel lumen before gas can be extracted from the adjacent vessel down the chain, or before it can be extracted from the gas dissolved in radially connected tissue. Therefore, what we might want to measure is the initial slope of *dP/dt*. There is experimental evidence supporting this finding, as the highest overall agreement between VCs based on the Pneumatron and a flow-centrifuge method was found after 15 s of gas extraction (Palighi *et al*., submitted).

Previous studies (Pereira *et al*. 2016, 2020; Zhang *et al*., 2018) computed *ΔP = P_i_ − P_t_*, where *P*_i_ is the initial pressure (at time zero) and *P*t is the pressure measured at time *t* = 60 s, 30 s, or 15 s. However, there is an inherent uncertainty in knowing time zero when the vacuum pump that draws down the pressure is turned off, and there is also an uncertainty of each pressure measurement *P_0_, P_1_… P_t_*; call this uncertainty ±δP_e_ and the time uncertainty ±δt. The combined uncertainty is then 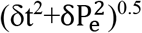. There could also be a transient period immediately after turning off the pump until the pressure is approximately equalized between the pressure sensor and the cut-open vessels. It could be argued that the best way to deal with this uncertainty, once you have optimized time and pressure measurements, is to perform a regression of *P_t_* versus *t*, in perhaps a 3-s time interval, and use the slope, *m*, of a linear regression to obtain the initial slope. Therefore, *PLC_Pn_* would be based on slope *m*.

Modeling results are often useful, but their precision may not correspond to the precision of real experimental results. Some insights can be gained from a brief look at real data. Currently, the time step used for Pneumatron measurements using a programmed Arduino-based system is 0.5 s. A cursory examination of a typical dataset taken during the dehydration of a *Eucalyptus* shoot with attached leaves revealed that the first few points after the pump was turned off followed a curvilinear trend during the first 2.5 s (Fig. 7a), but later appeared to be more linear. The slope versus time of dehydration showed a good trend for 3-s regression periods (seven-point regression from time 3 to 6 s, with measurements taken every 0.5 s) and for a 7-s regression period (13-point regression from time 3 to 10 s), but the slope for the longer regression period is less than that for the shorter period (Fig. 7b). The R^2^ values of the regressions showed marked differences during the 20-h dehydration experiment. This revealed that even shorter times for regressions rather than just pressure differences over 15 s might yield quite precise results.

**Figure 7.**
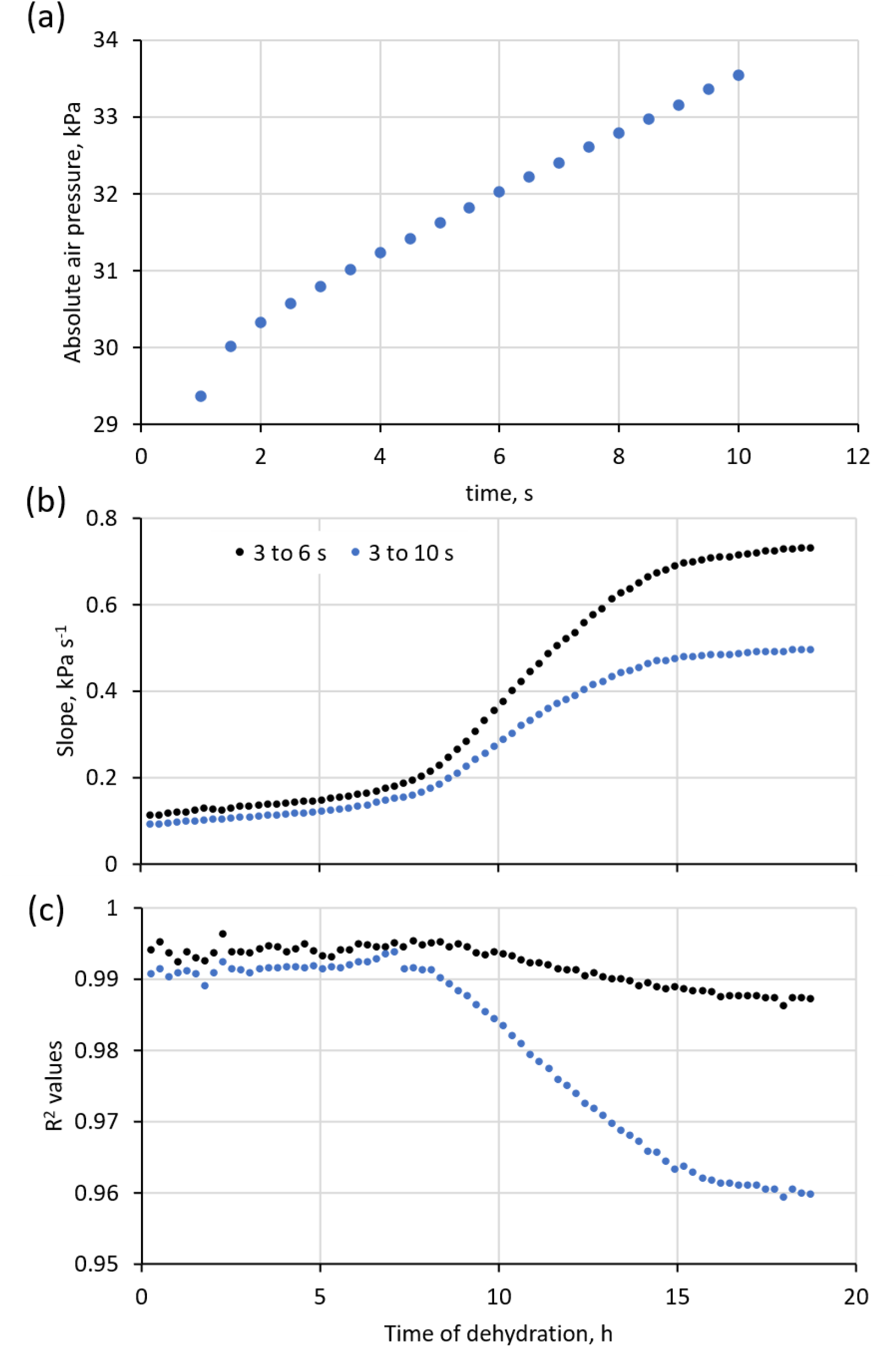
Analysis of experimental Pneumatron data collected during the dehydration of a *Eucalyptus camaldulensis* Dehnh. shoot with leaves. (a) Typical change in absolute pressure with time during shoot dehydration. (b) Slopes calculated for 3-s (black symbols) and 7-s periods (blue symbols) as affected by the dehydration time. Slopes were calculated after the initial 3 s, when there was a linear correlation between pressure and time, as shown in (a). (c) R^2^ values of slope values shown in (b) during dehydration time.

Using the UPPn model, we can also show that the *PLC_Pn_* values are closer to the real *PLC* values over the entire range of the VC (Table 3). While varying the number of vessels connected axially from 1 to 10, *PLC_Pn_* was 57.5% to 57.8% and, hence, depended somewhat on the number of vessels connected. However, when slope *m* was used, the *PLC_Pn_* was 52.3% and was much less dependent on the number of vessels in series. The absolute error of 2.4% in *PLC_Pn_* would cause a typical error of *T*50 by <30 kPa. The lack of dependence on the number of vessels in series is very fortunate, since we have no way of knowing how many vessels will be embolized in series as a function of *PLC*.

**Table 3.** This table used the default parameters in Table 2 in the Pneumatron Excel spreadsheet to compute the values shown below. The number of vessels in series (in the first column) connected axially was varied from 1 to 10. The 3-s interval used for these calculations was from t = 3 to 6 s.

| #vessels | First 15 s, $\Delta P$ (kPa) | | | PLC<sub>Pn</sub> | Slope (kPa s<sup>-1</sup>), 3 s | | | PLC<sub>Pn</sub> |
| --- | --- | --- | --- | --- | --- | --- | --- | --- |
| | $\Delta P_{\min}$ | $\Delta P_x$ | $\Delta P_{\max}$ | | $m_{\min}$ | $m_x$ | $m_{\max}$ | |
| 1 | 0.400 | 3.649 | 6.051 | 57.49% | 0.042 | 0.427 | 0.777 | 52.35% |
| 2 | 0.400 | 4.219 | 7.011 | 57.78% | 0.042 | 0.433 | 0.789 | 52.35% |
| 3 | 0.400 | 4.257 | 7.076 | 57.77% | 0.042 | 0.433 | 0.789 | 52.35% |
| 10 | 0.400 | 4.258 | 7.079 | 57.77% | 0.042 | 0.433 | 0.789 | 52.35% |

### Insights from a theoretical approach on plant pneumatics

Much more work on modeling of gas movement in stems seems to be merited by the encouraging results of this study. The tentative conclusion from the mathematical modeling of the biophysical process of gas movement in woody stems provides strong justification for the pneumatic method of measuring VCs and gas kinetics.

The main shortcoming of all mathematical models is that they cannot disprove experiments. This is because the validity of models always depends on the underlying assumptions made in the model. Thus, when a model provides confirmation of experimental results, it provides a theoretical basis for believing the experimental results. However, when results from well-designed and well-executed experiments disagree with a model, the model must always be presumed wrong.

If sometimes the pneumatic measurements produced a VC that readers found difficult to believe, it seems likely that it could be traced to methodological errors in the measurement of xylem water potential, and/or errors in the pneumatic measurements. For instance, incorrect estimation of the minimum and maximum amount of air discharge (Δ*P_min_*, and Δ*P_max_*, respectively), are well known to result in VCs that should be interpreted carefully (Chen *et al*., 2020; Sergent *et al*., 2020; Pereira *et al*., 2021). Users who are new to the Pneumatic method should pay special attention that stable measurements of Δ*P_min_* and Δ*P_max_* are obtained (Chen *et al*., 2020), which is easier when working with a Pneumatron than applying the manual pneumatic approach (Trabi *et al*., Submitted). The importance of which measurement values are considered as the functional starting and ending point also applies to hydraulic vulnerability curves (Choat *et al*., 2010; Jansen *et al*., 2015). Moreover, unstable values of Δ*P_min_* and Δ*P_max_* are much more likely to affect VCs than speculation about cracks in xylem (Chen *et al*., 2020), which would only affect pneumatic measurements if there would be direct cell wall openings to the intact conduits from which gas is extracted. Even if such xylem cracks would occur, it would result in a leakage that can easily be detected.

Although our model shows that pneumatic measurements are insensitive to vessel length (*L_v_*), another potential measuring error could result from the volume of the discharge tube. If the discharge volume is too large, the measuring error of the pressure sensor will be relatively large (see Fig. 4 in Jansen *et al*., 2020). We therefore recommend to adjust the volume of the discharge tube by determining the maximum volume of the gas that can be extracted from a completely dehydrated sample before conducting VC measurements (Pereira *et al*., 2020).

So far, the pneumatic method has not been successfully applied yet to Gymnosperms based on two species of Pinaceae (Zhang *et al*., 2018) and two species of Cupressaceae (Sergent *et al*., 2020). A possible explanation for a difference between angiosperms and gymnosperms could be aspiration of the pectin-rich torus in gymnosperms (Dute *et al*., 2015), preventing gas extraction from embolized tracheids. This could be tested by applying the Pneumatic method to gymnosperm species without a torus-margo pit membrane (Bauch *et al*., 1972), as suggested also by good agreement between pneumatic and hydraulic VCs of the vesselless angiosperm *Drimys brasiliensis* (Fig. 7 in Pereira *et al*., 2016).

Comparison of the Pneumatic with hydraulic and other non-hydraulic methods has provided strong agreement for a substantial number of angiosperm species and samples (Fig. S1 and Table S2; Pereira *et al*., 2016, 2020, 2021; Zhang *et al*., 2018, Sergent *et al*., 2020; Guan *et al*., 2021; Paligi *et al*., submitted). This agreement is especially strong when the above-mentioned caveats are considered. Therefore, we assert tentatively that there is a good theoretical and experimental basis for applying the Pneumatic method in research on plant water relations and embolism resistance.

## Acknowledgements

This research was funded by the National Natural Science Foundation of China (31770647) to D.M.Y, and a grant from the German Research Foundation (DFG, Deutsche Forschungsgemeinschaft, project nr. 410768178) to S.J. The authors acknowledge the São Paulo Research Foundation (FAPESP, Brazil) for a grant (M.T.T. and R.V.R., grant #2019/24519-1) and fellowship (L.P. and R.V.R., grant #2017/14075-3). R.V.R. is a fellow of the National Council for Scientific and Technological Development (CNPq, Brazil).

## Supporting information

Additional supporting information may be found in the online version of this article.

**Fig. S1** Comparison of embolism resistance measured with the Pneumatic method and other methods.

**Table S1** Excel spreadsheet of the UPPn model to simulate gas kinetics.

**Table S2** Published values of embolism resistance estimated with the Pneumatic and other methods.

**Note S1** Description of the basic layout of the UPPn model as shown in the Excel spreadsheet.

