## Supplementary Material for "A Unit Pipe Pneumatic model to simulate gas kinetics during measurements of embolism in excised angiosperm xylem"

The following Supporting Information is available for this article:

**Fig. S1** Comparison of embolism resistance measured with the Pneumatic method and other methods.

**Table S1** Excel spreadsheet of the UPPn model to simulate gas kinetics.

**Table S2** Published values of embolism resistance estimated with the Pneumatic and other methods.

**Notes S1** Description of the basic layout of the UPPn model as shown in the Excel spreadsheet.

**Fig. S1** $\Psi_{50}$ values estimated with the Pneumatic method as compared to other methods based on literature (N = 57 specimens, including 51 species), grouped by reference (a), or alternative method to measure embolism resistance (b). Only angiosperms were considered; outliers due to measurement errors were not removed (see Pereira *et al*. (2021), and the discussion in the main text). All data are provided in Table S2. The black line represents the 1:1 reference line.

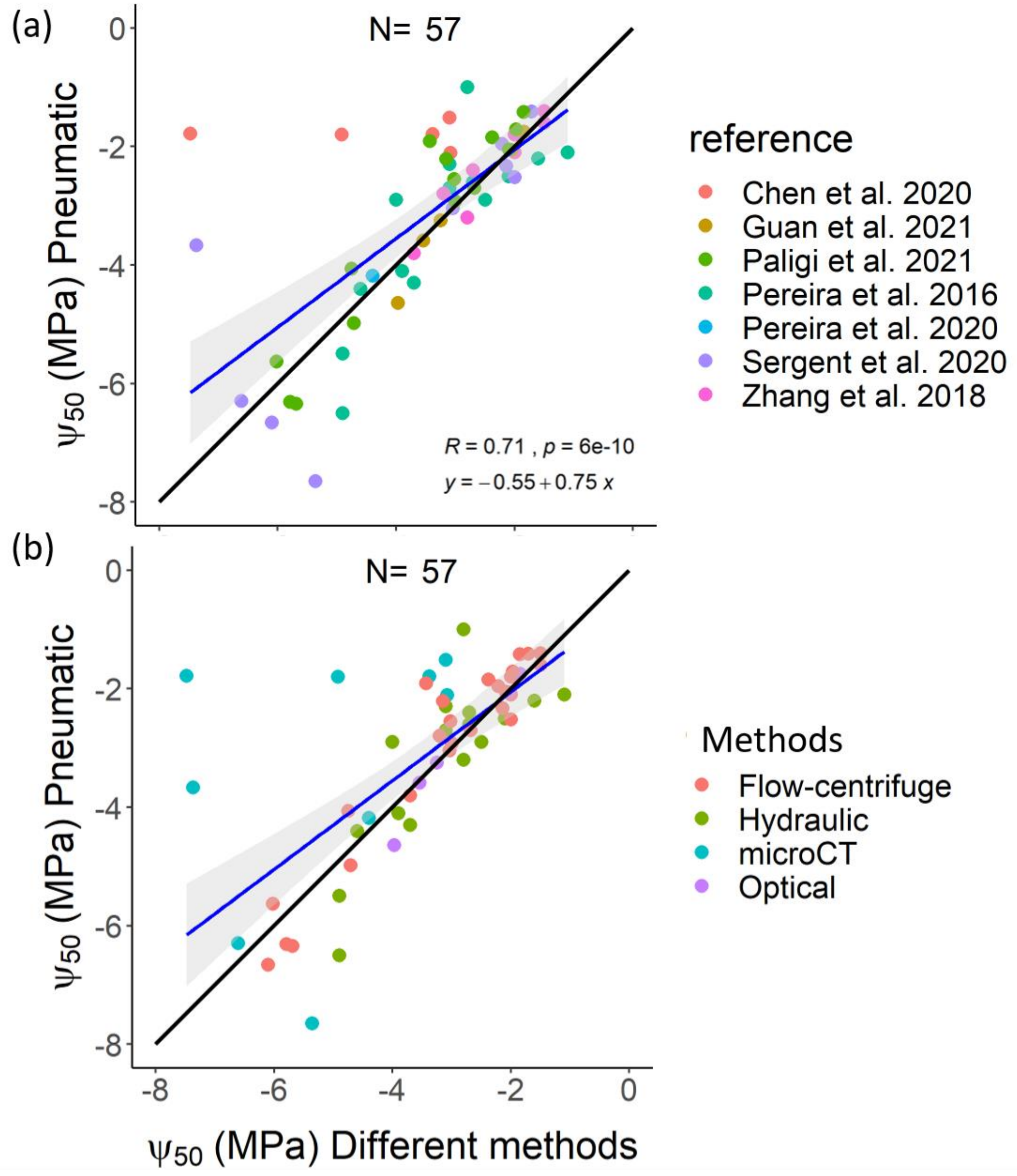

**Table S2** Published $\Psi_{50}$ values estimated with the Pneumatic and alternative methods. Species are listed alphabetically, with reference to the plant organ studied, maximum vessel length (MLV), the alternative method applied to measure embolism resistance, and the reference. The column "Similar plant material?" indicates whether or not similar samples from the same plant were used.

| Species | Organ | MVL (m) | $\Psi_{50}$ pneumatic | Alternative method | $\Psi_{50}$ alternative | Reference | Similar plant material? |
|---|---|---|---|---|---|---|---|
| *Alnus glutinosa* | stem | | -1.6 | Flow-centrifuge | -1.5 | Zhang et al. 2018 | yes |
| *Austrocedrus chilensis* | stem | Tracheids | -2.05 | Flow-centrifuge | -4.02 | Sergent et al. 2020 | yes |
| *Betula pendula* | stem | | -1.8 | Flow-centrifuge | -2 | Zhang et al. 2018 | yes |
| *Betula pendula* | stem | | -1.85 | Flow-centrifuge | -2.38 | Paligi et al. submitted | yes |
| *Betula utilis* | stem | | -2.05 | Flow-centrifuge | -2.09 | Paligi et al. submitted | yes |
| *Calyptranthes brasiliensis* | stem | 0.474 | -5.5 | Hydraulic | -4.9 | Pereira et al. 2016 | yes |
| *Carpinus betulus* | stem | | -3.8 | Flow-centrifuge | -3.7 | Zhang et al. 2018 | yes |
| *Carpinus betulus* | leaf | | -3.25 | Optical | -3.25 | Guan et al. 2021 | yes |
| *Carpinus betulus* | stem | | -4.06 | Flow-centrifuge | -4.75 | Paligi et al. submitted | yes |
| *Cedrus deodara* | stem | Tracheids | -3.72 | Flow-centrifuge | -6.01 | Sergent et al. 2020 | yes |
| *Citrus sinensis* | stem | 0.362 | -4.3 | Hydraulic | -3.7 | Pereira et al. 2016 | yes |
| *Coffea arabica* | stem | 0.4 | -2.9 | Hydraulic | -2.5 | Pereira et al. 2016 | yes |
| *Combretum griffithii* | stem | 1.41 | -1.51 | MicroCT | -3.1 | Chen et al. 2020 | yes |
| *Combretum yunnanense* | stem | 1.26 | -1.8 | MicroCT | -4.92 | Chen et al. 2020 | yes |
| *Corylus avellana* | stem | | -2.1 | Flow-centrifuge | -2 | Zhang et al. 2018 | yes |
| *Crataegus persimilis* | stem | | -5.63 | Flow-centrifuge | -6.02 | Paligi et al. submitted | yes |
| *Cupressus sempervirens* | stem | Tracheids | -8 | Hydraulic | -10.4 | Pereira et al. 2016 | no |
| *Drimys brasiliensis* | stem | Tracheids | -2.2 | Hydraulic | -1.6 | Pereira et al. 2016 | yes |
| *Embothrium coccineum* | stem | 0.11 | -1.41 | Flow-centrifuge | -1.71 | Sergent et al. 2020 | yes |
| *Erisma uncinatum* | stem | 0.41 | -2.1 | Hydraulic | -1.1 | Pereira et al. 2016 | yes |
| *Eucalyptus camaldulensis* | stem | 0.46 | -4.1 | Hydraulic | -3.9 | Pereira et al. 2016 | yes |

| | | | | | | | |
|---|---|---|---|---|---|---|---|
| *Fagus sylvatica* | stem | | -2.8 | Flow-centrifuge | -3.2 | Zhang et al. 2018 | yes |
| *Fagus sylvatica* | leaf | | -2.06 | Optical | -2.05 | Guan et al. 2021 | yes |
| *Fraxinus excelsior* | stem | | -2.4 | Hydraulic | -2.7 | Zhang et al. 2018 | yes |
| *Hymenaea courbaril* | stem | 0.15 | -1 | Hydraulic | -2.8 | Pereira et al. 2016 | no |
| *Lagerstroemia tomentosa* | stem | 0.99 | -2.11 | MicroCT | -3.08 | Chen et al. 2020 | yes |
| *Lasiococca comberi* | stem | 0.39 | -1.78 | MicroCT | -7.48 | Chen et al. 2020 | yes |
| *Laurus nobilis* | stem | 0.9 | -3.67 | MicroCT | -7.37 | Sergent et al. 2020 | no |
| *Liriodendron tulipifera* | stem | | -1.4 | Flow-centrifuge | -1.5 | Zhang et al. 2018 | yes |
| *Liriodendron tulipifera* | leaf | | -1.75 | Optical | -1.84 | Guan et al. 2021 | yes |
| *Lomatia hirsuta* | stem | 0.12 | -1.96 | Flow-centrifuge | -2.21 | Sergent et al. 2020 | yes |
| *Maytenus boaria* | stem | 0.22 | -6.66 | Flow-centrifuge | -6.1 | Sergent et al. 2020 | yes |
| *Miconia lepidota* | stem | 0.25 | -4.4 | Hydraulic | -4.6 | Pereira et al. 2016 | yes |
| *Microcos paniculate* | stem | 0.37 | -1.79 | MicroCT | -3.38 | Chen et al. 2020 | yes |
| *Myrceugenia acutata* | stem | 0.662 | -6.5 | Hydraulic | -4.9 | Pereira et al. 2016 | yes |
| *Nothofagus antarctica* | stem | 0.19 | -2.33 | Flow-centrifuge | -2.14 | Sergent et al. 2020 | yes |
| *Nothofagus pumilio* | stem | 0.16 | -2.52 | Flow-centrifuge | -2 | Sergent et al. 2020 | yes |
| *Olea europaea* | stem | 0.293 | -2.9 | Hydraulic | -4 | Pereira et al. 2016 | no |
| *Olea europaea* | stem | 0.83 | -7.65 | MicroCT | -5.36 | Sergent et al. 2020 | no |
| *Olea europaea* | stem | | -4.18 | MicroCT | -4.4 | Pereira et al. 2021 | no |
| *Ostrya carpinifolia* | stem | | -4.98 | Flow-centrifuge | -4.71 | Paligi et al. submitted | yes |
| *Pinus pinaster* | stem | Tracheids | -2.8 | Flow-centrifuge | -3.7 | Zhang et al. 2018 | yes |
| *Pinus sylvestris* | stem | Tracheids | -2 | Flow-centrifuge | -3.2 | Zhang et al. 2018 | yes |
| *Platanus acerifolia* | stem | | -1.42 | Flow-centrifuge | -1.85 | Paligi et al. submitted | yes |
| *Platanus orientalis* | stem | | -1.71 | Flow-centrifuge | -1.97 | Paligi et al. submitted | yes |
| *Populus nigra* | stem | 0.163 | -2.6 | Hydraulic | -2.7 | Pereira et al. 2016 | no |
| *Prunus avium* | leaf | | -4.64 | Optical | -3.97 | Guan et al. 2021 | yes |
| *Pyrus calleryana* | stem | | -6.31 | Flow-centrifuge | -5.79 | Paligi et al. submitted | yes |
| *Quercus ilex* | stem | 0.9 | -6.3 | MicroCT | -6.61 | Sergent et al. 2020 | no |
| *Quercus petraea* | leaf | | -3.59 | Optical | -3.54 | Guan et al. 2021 | yes |
| *Quercus robur* | stem | | -3.2 | Hydraulic | -2.8 | Zhang et al. 2018 | yes |
| *Schinus patagonicus* | stem | 0.14 | -3.04 | Flow-centrifuge | -3.04 | Sergent et al. 2020 | yes |

| | | | | | | | |
|---|---|---|---|---|---|---|---|
| *Schinus terebinthifolius* | stem | 0.187 | -2.3 | Hydraulic | -3.1 | Pereira et al. 2016 | yes |
| *Schinus terebinthifolius* | stem | 0.254 | -2.3 | Hydraulic | -3.1 | Pereira et al. 2016 | yes |
| *Schinus terebinthifolius* | stem | 0.254 | -2.7 | Hydraulic | -3.1 | Pereira et al. 2016 | yes |
| *Sorbus latifolia* | stem | | -6.34 | Flow-centrifuge | -5.69 | Paligi et al. submitted | yes |
| *Thuja plicata* | stem | Tracheids | -7.6 | Hydraulic | -5 | Pereira et al. 2016 | no |
| *Tilia cordata* | stem | | -2.55 | Flow-centrifuge | -3.02 | Paligi et al. submitted | yes |
| *Tilia cordata* | stem | | -2.21 | Flow-centrifuge | -3.15 | Paligi et al. submitted | yes |
| *Tilia japonica* | stem | | -2.92 | Flow-centrifuge | -3 | Paligi et al. submitted | yes |
| *Tilia platyphyllos* | stem | | -2.7 | Flow-centrifuge | -2.68 | Paligi et al. submitted | yes |
| *Tilia platyphyllos* | stem | | -1.91 | Flow-centrifuge | -3.43 | Paligi et al. submitted | yes |
| *Weinmannia organensis* | stem | 0.497 | -2.5 | Hydraulic | -2.1 | Pereira et al. 2016 | yes |

**Notes S1** Basic layout of the UPPn Excel spreadsheet

For the Excel spreadsheet, the values in yellow must be entered, and the values in white are computed from the yellow values (Table 2). The first two values are physical constants that should not be altered. The next two values are diffusion coefficients for gases in lignified wood ($D_{gw}$) (Sorz and Hietz 2006) and in wet pit membranes ($D_g$), which might optionally be reduced by 20% to allow for the pit membrane having solid cellulose fibers. The following three values are all available from the literature (Zanne et al. 2010, Morris et al. 2016) or can be measured anatomically: (1) the fraction of vessel walls in common between vessels is available in some studies (Sperry et al. 2005, 2006), (2) the percentage of stem cross section that is vessel, and (3) vessel diameter. The fraction of pits in the pit field varies over a narrow range from 0.4 to 0.6 (Lens et al. 2011, Scholz & Jansen 2013). The thickness of pit membranes varies from 0.2 to 1.3 μm (Li et al. 2016, Kaack et al. 2019). *dt* is a step time interval that must be small enough for the computational results to be stable. This value is determined by trial and error and is impacted by all the other parameters. The last two values are the volume of tubing connected to the pressure transducer and the diameter of the wood. These values are used to compute the number of vessels and the external volume per vessel, because the UPPn model needs to scale the volume where pressure is measured in a single vessel series (unit pipe). In the numerical simulation, there are only two types of rate constants that must be calculated, and they depend on the values above. One type of rate constant is for the radial diffusion, and the other is for axial diffusion, as noted in the previous section.

The basic layout of the Excel spreadsheet is as follows: There are 3070 rows of data, and columns extend from A to DS. Row 1 is sufficiently widened to contain many output graphs of gas concentration in the embolized vessels and water-filled tissue around the vessels. Every cell with a computed value can be plotted in these graphs. Rows 2–28 are used to present the basic equations in text boxes or user-entered values for calculations. Rows 29–37 are mass conservation checks. It is very important to check that many thousands of equations embedded in the cells are correct. If they are correct, then the mass (moles) will be preserved; that is, the loss of moles from the embolized tissue should equal the gain of moles in the volume space where pressure is measured with the Pneumatron. Rows 43–3070 contain computed outputs in concentration units (mol $m^{-3}$) in liquid and gas phases, consistent with the requirements of Henry's law. Some columns contain *dn* moles, that is, the number of moles in a time interval, *dt*, passing between vessels by diffusion through pit membranes. Columns CX–DS starting at row 46 are gas phase concentrations, $C_g$, converted to pressure, $P$, with the ideal gas law $P = C_g RT = nRT/V$, and the results in graphical form.

For readers who want to use the Excel spreadsheet for their own computations, the publisher's download site for supplemental materials will contain the Excel spreadsheet and a pdf file with more details. Readers can also experiment with different formulas in Excel cells if desired. The mass balance equations are useful and should not be altered. Mass balance (conservation of mass) is necessary but not sufficient proof that the model is correct. Errors in equations can cause a violation of mass balance.
